# A Zebrafish Platform to Model Human *SCN2A* and *SCN8A* Epilepsy and Evaluate Anti-Seizure Medications

**DOI:** 10.64898/2026.06.21.733632

**Authors:** Patrick C Milder, Theodore R Cummins, James A Marrs

## Abstract

Many patients with epilepsy have inadequate seizure control using current anti-seizure medications (ASMs), illustrating the need for new treatments. Genetic epilepsy syndromes like pathogenic variants in voltage gated sodium channel *SCN2A* and *SCN8A* are often poorly controlled by current medications, highlighting the need for better models. Voltage gated sodium channel pathogenic variants that induce epilepsy are often gain-of-function, producing hyperexcitability. We established a fast and precise zebrafish seizure assay using mRNA overexpression of *SCN2A* and *SCN8A* variants, which allows rapid screening of both variants and ASMs. These short-term genetic seizure models are assayed in 3 days postfertilization (dpf) larvae. Pathogenic variants of *SCN2A* and *SCN8A* produced sporadic seizure behavior. We tested human *SCN2A* R1882Q, *SCN2A* R853Q and *SCN8A* R1872Q pathogenic variants that were identified in epilepsy syndrome patients. These models were used to evaluate the efficacy of 3 ASMs: Topiramate, GS967 and PF-04856264. All 3 epilepsy-associated variants increased seizure activity, and the ASMs significantly decreased this seizure activity. This mRNA overexpression assay successfully evaluates seizure activity induced by variants in voltage gated sodium channel genes and examines ASM efficacy in patient specific pathogenic variants.

**Summary:** Genetic epilepsy syndromes have devastating consequences for patients. A zebrafish method is presented that analyzes pathogenic epilepsy variants in human voltage gated sodium channels (Na_v_1.2 and Na_v_1.6) and tests efficacy of anti-seizure medications (ASMs). This method provides a rapid platform for evaluating variants and medications.

## Introduction

Epilepsy is a disorder characterized by recurrent seizures, which are often caused by an imbalance between inhibition and excitation (Scharfman, 2007). Approximately thirty percent of patients with epilepsy are refractory to currently available treatments. Thus, there is a critical need to identify and screen anti-seizure medications (ASMs). The voltage gated sodium channels (Na_v_s) are central to neuron excitability. This study specifically focuses on Na_v_1.2 and Na_v_1.6 found at the pre- and post-synaptic membranes (Pan and Cummins, 2020). Na_v_1.2 and Na_v_1.6 subunits contribute to action potential generation. Genetic variants of Na_v_1.2 and Na_v_1.6 affect channel gating and may lead to hyperexcitability associated with seizures (Meisler et al., 2016; Patel et al., 2016; Sanders et al., 2018).

*SCN2A* codes for Na_v_1.2 and is initially expressed at early stages of neural development (Catterall, 2014). As development continues, Na_v_1.2 axonal expression becomes more restricted to unmyelinated axons, and Na_v_1.6 is prominently located at the nodes of Ranvier in myelinated axons (Van Wart and Matthews, 2006), replacing Na_v_1.2 in many developing axons. This process is also seen across the peripheral nervous system (Van Wart and Matthews, 2006). Both Na_v_1.2 and Na_v_1.6 can be found in axon initial segments of excitatory neurons (Katz et al., 2018).

In the zebrafish, a homolog of human *SCN2A* is not defined, but *scn8aa* and *scn8ab* are homologous to human *SCN8A*. Subcellular protein distributions are not known in the zebrafish because specific antibody reagents are not yet available. Zebrafish has two Na_v_1.6 encoding genes, *scn8aa* and *scn8ab*. The *scn8aa* mRNA expression in the central nervous system was detected in the brain, the otic vesicle, spinal sensory Rohon-Beard cells and spinal motor neurons (Novak et al., 2006). Expression of *scn8ab* is much more restricted (Novak et al., 2006).

Pathogenic Na_v_ variants can cause epilepsy (Estacion et al., 2014). *SCN2A* pathogenic variants cause severe phenotypes such as West Syndrome (Epi et al., 2013), Ohtahara Syndrome (Allen et al., 2016), and infantile spasms (Ogiwara et al., 2009). There have been over 150 frequently occurring *SCN2A* variants associated with epileptic phenotypes (Wolff et al., 2017; Mason et al., 2019). Two commonly documented variants in *SCN2A* associated with epilepsy are R1882Q and R853Q (Carvill et al., 2013; Epi et al., 2013; Nakamura et al., 2013; Howell et al., 2015; Samanta and Ramakrishnaiah, 2015; Kobayashi et al., 2016; Li et al., 2016; Trump et al., 2016; Wolff et al., 2017). *SCN2A* R1882Q is a gain-of-function variant, causing increased action potential frequency (Li et al., 2021), which lead to seizures and epilepsy. The *SCN2A* R853Q variant has been identified in multiple patients with epilepsy, but the functional consequences are less clear. Studies have suggested that R853Q might be a loss of function variant (Berecki et al., 2018; Ganguly et al., 2021), but recent work showed that R853Q produces robust gating pore currents suggesting it might have gain of function attributes (Mason et al., 2019). This highlights the need for additional functional screens for sodium channel variants, like the *SCN2A* variant R853Q, to better determine their impact on overall activity and seizure generation.

*SCN8A* encodes Na_v_1.6, which regulates and maintains neuronal excitability in the central nervous system (Do and Bean, 2004). Na_v_1.6 channels are expressed in excitatory neurons, concentrated in the axon initial segment and nodes of Ranvier, where action potentials are initiated and propagated (Hu et al., 2009). Loss-of-function SCN8A variants often produce cognitive defects, developmental delay, and autism symptoms without epilepsy (Johannesen et al., 2022). There are other cases where loss-of-function variants produce epilepsy symptoms, often with late onset or absence seizures (Johannesen et al., 2022). Gain-of-function SCN8A variants most commonly produce early onset epilepsy, including developmental and epileptic encephalopathy (Veeramah et al., 2012; de Kovel et al., 2014; Estacion et al., 2014; Blanchard et al., 2015; Wagnon et al., 2015; Anand et al., 2016; Barker et al., 2016; Patel et al., 2016). Over 200 *SCN8A* variants show epileptic phenotypes that include epileptic encephalopathy, refractory epilepsy and other debilitating conditions (Johannesen et al., 2022). We investigated the *SCN8A* gain-of-function variant, R1872Q, which was chosen because the amino acid change occurs at the comparable position in the Na_v_1.6 channel as the Na_v_1.2 channel variant R1882Q. The *SCN8A* R1872Q variant causes premature activation and increased resurgent currents in the Na_v_1.6 channel (Pan and Cummins, 2020).

Here, a zebrafish platform was developed to evaluate human *SCN2A* and *SCN8A* variants that are associated with epilepsy. This experimental paradigm allowed us to test if specific variants have gain of function phenotype and test different ASMs against these patient specific variants. Testing variants found in patients could support translational and personalized medicine research approaches. Epilepsy variants occurring in the voltage-gated sodium channel alpha-subunits were evaluated. For comparison, the human *SCN2A* R937C loss-of-function variant was tested, as this variant is associated with intellectual disability and autism spectrum disorder, but does not induce seizures in patients (Berecki et al., 2022). We propose this zebrafish platform will permit screening of many variants and ASMs, evaluating the impact of variants on overall channel function and drug efficacy on patient specific Na_v_ variants.

## Methods

### Zebrafish husbandry

All experiments and procedures followed the Indiana University Policy on Animal Care and Use guidelines and were approved by the IUPUI School of Science Institutional Animal Care and Use Committee, and all authors comply with ARRIVE guidelines. Zebrafish (*Danio rerio*) AB strain were raised and maintained under standard laboratory conditions (Westerfield, 2000). Fertilized eggs were collected from mating chambers and rinsed with embryo medium (EM). Zebrafish larva and embryos were maintained at 28.5°C, on a 14/10-hour light/dark cycle under standard conditions.

### Microinjection

Injection needles for mRNA injections were pulled from borosilicate glass capillary tubes containing filaments using a Sutter P1000 Flaming/Brown micropipette puller (Novato CA). Embryos were injected with the WPI PV280 (World Precision Instruments, Sarasota FL) through the chorion into the cell compartment at the one-cell stage. Plasmids for the respective cDNA sequences: human *SCN2A*-wt; *SCN2A*-R1882Q; *SCN2A*-R853Q; *SCN2A*-R937C; *SCN8A*-wt; and *SCN8A*-R1872Q were previously generated in codon optimized cDNA and cloned into the pcDNA3.1+ vector (Mason et al., 2019; Pan and Cummins, 2020). For synthesis of mRNAs, plasmids were cut with *XbaI* and synthesized with the mMessage Machine SP6 Kit (Life Technologies/Ambion, Darmstadt, Germany) according to the manufacturer’s instructions. Injected mRNA concentration was 400 ng/µL for *SCN2A* mRNAs and 800 ng/µL for *SCN8A* mRNAs.

### Experimental design and drug treatment

At 9 am, 3 dpf zebrafish were moved to individual wells of a 96-well plate and placed in 250 µL EM. In the *SCN2A* behavioral assays, five injected groups were used. The injection groups were as follows: H_2_O injected, *SCN2A-wt* mRNA injected, R937C mRNA injected, R1882Q mRNA injected, and R853Q mRNA injected. These five groups were treated with embryo medium or an anti-seizure compound. This resulted in ten treatment groups per plate with nine larvae per treatment group. Four replicate plates were done using either topiramate (TPR), GS967, and PF-04856264, making the total number of individuals was 36 per treatment group. In the *SCN8A* behavioral assays, three injection groups were used. The injection groups were as follows: H_2_O injected, *SCN8A-wt* mRNA injected, and R1872Q mRNA injected. These three groups were treated with embryo medium or an anti-seizure compound. This resulted in six treatment groups per plate with twelve larvae per treatment group. Three replicate plates were done using either TPR, GS967 and PF-04856264, making the total number of individuals 36 per treatment group.

Larvae were raised together in Petri plates with EM to control for incubation conditions, and all injection groups were assayed together on each 96-well plate to control for any plate variation. Injected larvae did not show abnormal morphology. Any larvae with abnormal morphology were eliminated from the experiment. After 1 pm, 3 dpf larvae were allowed to habituate for 30 minutes in the light at room temperature. After habituation, 250 µL of embryo medium or 250 µL of the 2x concentration of ASM compound solution was added. This method was used in all experiments.

Of the three ASMs (TPR, GS967, and PF-04856264) tested, PF-04856264 had not been tested in zebrafish, and thus, it was tested in at a range of concentrations: 0.1 µM, 1 µM, 10 µM and 100 µM. The 10 µM concentration consistently reduced seizure activity of the *SCN2A* R1882Q and *SCN2A* R853Q as compared with untreated groups in our pilot experiments. Based on previous work, 200 µM TPR (Afrikanova et al., 2013; Milder et al., 2022) and 0.05 µM GS967 (Milder et al., 2022) were used. Dimethyl sulfoxide (DMSO) was used to dissolve the ASMs and diluted to 0.1% DMSO for all treatments. Control experiments tested and compared to the epileptic variant mRNA injected groups with EM to 0.1% DMSO, which showed no significant difference.

### Movement tracking system

After 1 pm, larvae were moved to an automated tracking device, the ZebraBox apparatus, which tracks larvae movement and collects data. The large movement count best distinguished the rapid seizure movements that was quantified using ZebraLab software, which is defined as the number of movement events over a speed of 8 mm/s. Ten-minute integration periods were used. The first 30-minutes is an acclimation period in the dark. The acclimation period is not measured. Locomotion was tracked and measured over a 90-minute assay. After the 30-minute acclimation period, the fish are exposed to light for 10-minutes. Then, the light is turned off for 10-minutes. This protocol has alternating light for 10-minutes (five times) and dark for 10-minutes (four times) during the 90-minute assay.

### Statistical analysis

Locomotor behavior data between the replicate assays for injection groups and an ASM were averaged at each 10-minute period and compared by two-way ANOVA and Tukey’s post-hoc comparisons (GraphPad Prism software version 8.2, Boston MA). The pooled large counts of each treatment group over the 90-minute light-dark assay were compared using one-way ANOVA followed by a Tukey’s post hoc test (GraphPad Prism software version 8.2). Embryo medium was used as the control group for ASM treatment, and H_2_O injected groups were used as a vehicle control for mRNA injections.

## Supplementary Material

Tables showing statistics for data figures are shown in Supplementary Tables 1-13. Representative tracings of swimming behaviors for mRNA injected zebrafish illustrate the seizure behavior. Videos showing representative swimming behaviors of mRNA injected zebrafish larvae illustrate seizure behaviors (rapid circular swimming and tonic seizure or “freezing” behavior).

## Results

### Pathogenic *SCN2A* variant expression in larvae induces seizure behavior

We hypothesized that expressing gain of function pathogenic sodium channel genetic variants would induce seizure behavior in zebrafish larvae, providing a potential platform to study genetic seizure disorders. First, the effects of the *SCN2A* variant R1882Q was tested, as multiple studies have identified this variant as having strong gain of function (Berecki et al., 2018; Mason et al., 2019; Thompson et al., 2023). Increasing mRNA concentrations were injected into 1-cell embryos, and larvae behavior was measured in 3 dpf zebrafish. Rapid movements (>8mm/s) were counted as a surrogate for seizure activity. The behavior assay took place over a 120-minute duration, involving a 30-minute acclimation period in the dark, followed by five cycles of 10 minutes in the light and four cycles of 10 minutes in the dark. It is interesting that the light/dark movement response in zebrafish larvae develops on or after 5 dpf (Orger and de Polavieja, 2017), but light/dark promoted seizure activity in our experiments at 3 dpf. Only data during this 90-minute period of alternating light and dark were used in the analysis. Overall, increased movement was compared to the vehicle (H_2_O) injected group. Injection of 400 ng/μL mRNA produced the most consistent behavior response. Large count was found to be the most reliable indicator of increased activity caused by voltage gated sodium channel variant expression. The ZebraBox measures swimming speeds (inactive, 0-4 mm/second; small, 4-8 mm/second; and large, >8 mm/sec), giving counts (numbers of times larvae swim at those speeds), distance (total distances larvae swim at those speeds) and time (amount of time the larvae swim at those speeds). A schematic diagram of experimental procedures is shown in Fig. 1. Behavior responses are seen in tracings of swimming (Supplemental data, Fig S1) from the ZebraBox: inactive movement are black lines, small movement are green lines and large movement are red lines. Seizure behavior is seen in video sequences of *hSCN2A* (Supplemental data, Movie S1) and *hSCN8A* (Supplemental data, Movie S2) injected larvae, showing characteristic seizure activity that include rapid, circle swimming followed by freezing and loss of buoyant upright posture. If overexpression of the channels is due to ectopic expression in tissues outside the nervous system, then it is useful to note that wild type channels do not induce seizure behavior.

**Figure 1.**
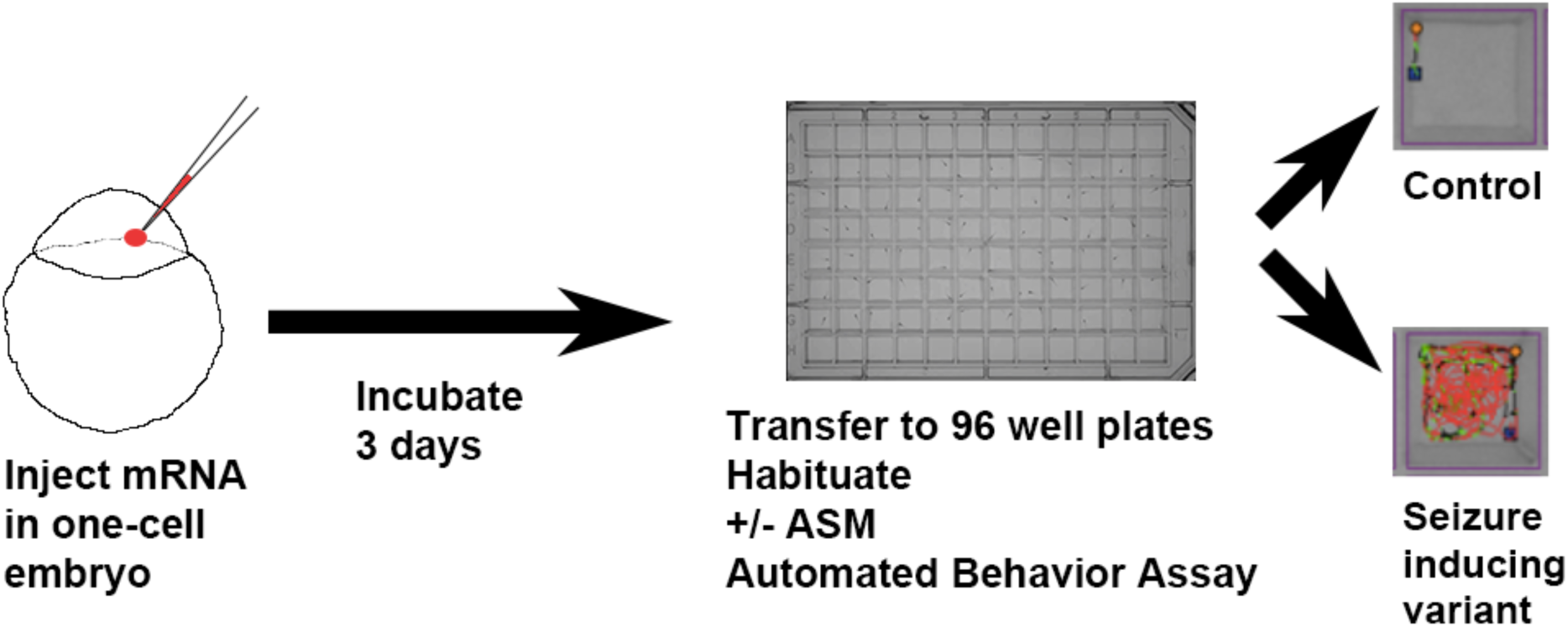
Schematic illustration of behavioral assay voltage gated sodium channel mRNA injected zebrafish larvae. One- or two-cell stage embryos are injected with mRNAs to express voltage gated sodium channel variants. Embryos develop into 3 dpf larvae, which are placed in 96-well plates for behavior tracking. After habituation in assay plate, some larvae are treated with ASMs. Automated behavior tracking is performed. Tracings of swimming behavior are shown for control and seizure inducing mRNA variants (red lines show large/rapid movements).

The wild type (*SCN2A-*wt) mRNA injection did not induce seizure behavior (Figs 2A, 3A & 4A). The R1882Q *SCN2A* pathologic variant mRNA overexpression produced statistically significant increases in seizure behavior in all assays (Figs 2B, 3B & 4B). A strong loss of function *SCN2A* variant that does not induce seizures in humans, R937C, (Ben-Shalom et al., 2017) was compared to evaluate assay specificity. There was no significant difference in large count between the H_2_O injected, *SCN2A*-wt or R937C groups in any of our behavioral assays (Figs 2A, 3A & 4A). Finally, the R853Q *SCN2A* pathologic variant, which is associated with seizures but has mixed loss of function and gain of function effects on channel currents was tested. R853Q mRNA overexpression produced statistically significant increases in seizure behavior in all assays (Figs 2C, 3C & 4C). The total large count summed over the 90-minute assay was compared, showing that the R1882Q and R853Q groups produced significantly higher total large counts as compared to the H_2_O injected larvae (Fig 2D, 3D & 4D). These data show that R853Q has an overall gain of function effect compared to wild type Nav1.2 channels.

**Figure 2.**
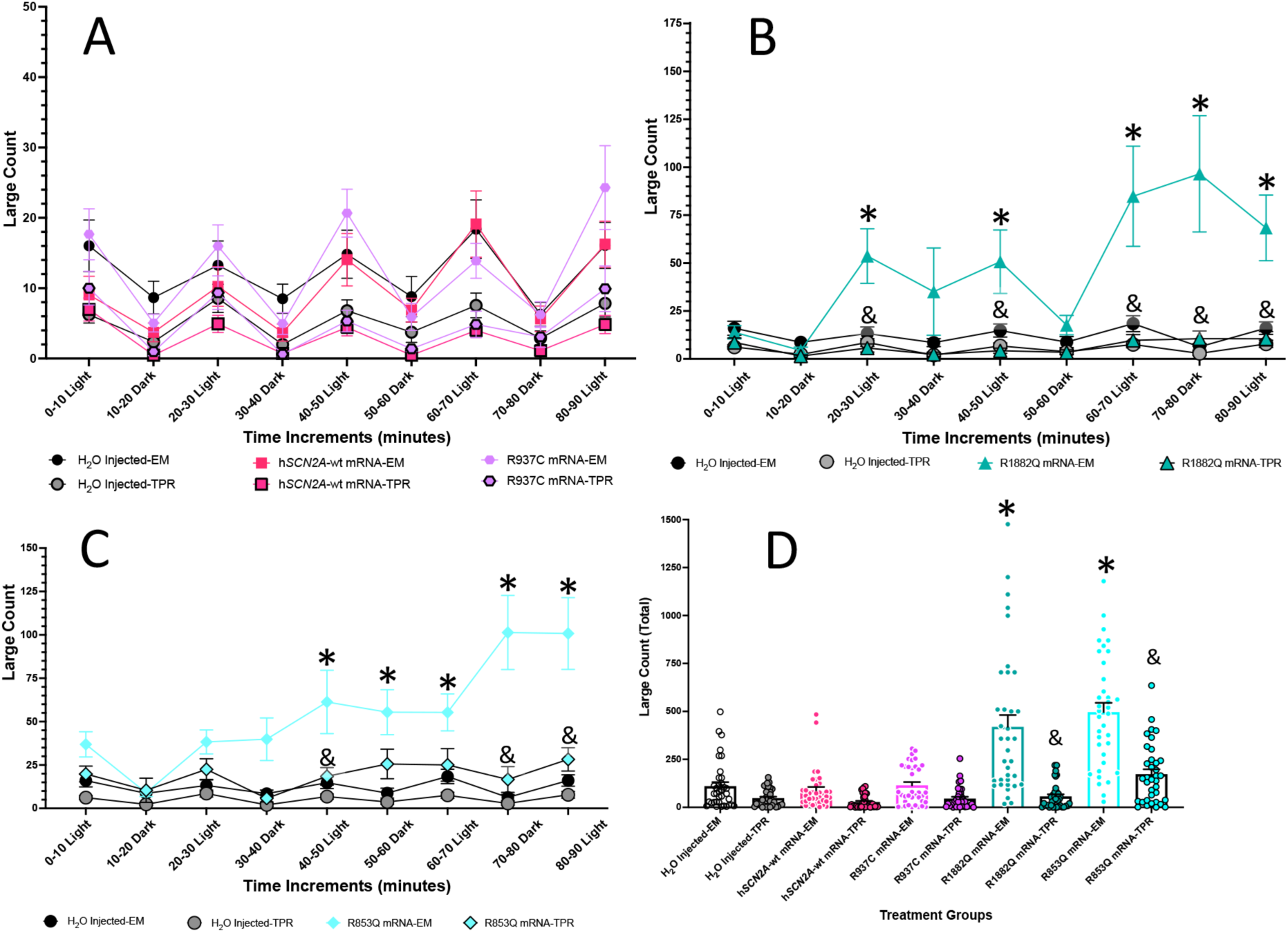
Behavioral profile and effects of topiramate on 3 dpf h*SCN2A* injected zebrafish larvae. (A-C) The average large count, number of large movement events (y-axis) of larvae is plotted against time, alternating 10-minute cycles of light and dark (x-axis) during the tracking experiment. The treatment groups are listed below the x-axis, showing the shape and color of the graph points and lines. Embryo medium (EM) is compared with ASM treatment. (A) Graph compares TPR treatment of H_2_O injected, wild type human *SCN2A* mRNA injected, and non-seizure inducing human *SCN2A* R937C mRNA injected with untreated groups. (B) Graph compares TPR treatment of H_2_O injected and seizure inducing human *SCN2A* R1882Q mRNA injected with untreated groups. (C) Graph compares TPR treatment of H_2_O injected and seizure inducing human *SCN2A* R853Q mRNA injected with untreated groups. (D) A histogram representing the total large count (y-axis) summed over the 90-minute assay shows the various treatment groups. The respective treatment groups are listed below the x-axis. Statistical significance (two-way ANOVA, Tukey’s post-hoc) are marked (*=p<0.05) to indicate comparison to control, H_2_O -injected embryos. Statistical significance (two-way ANOVA, Tukey’s post-hoc) are marked (&=p<0.05) to indicate comparison to h*SCN2A*-variant vs h*SCN2A*-variant + TPR. Standard error of the mean is shown for each. n = 36 fish/treatment group. Statistics are found in the Supplementary Table S1-S2.

**Figure 3.**
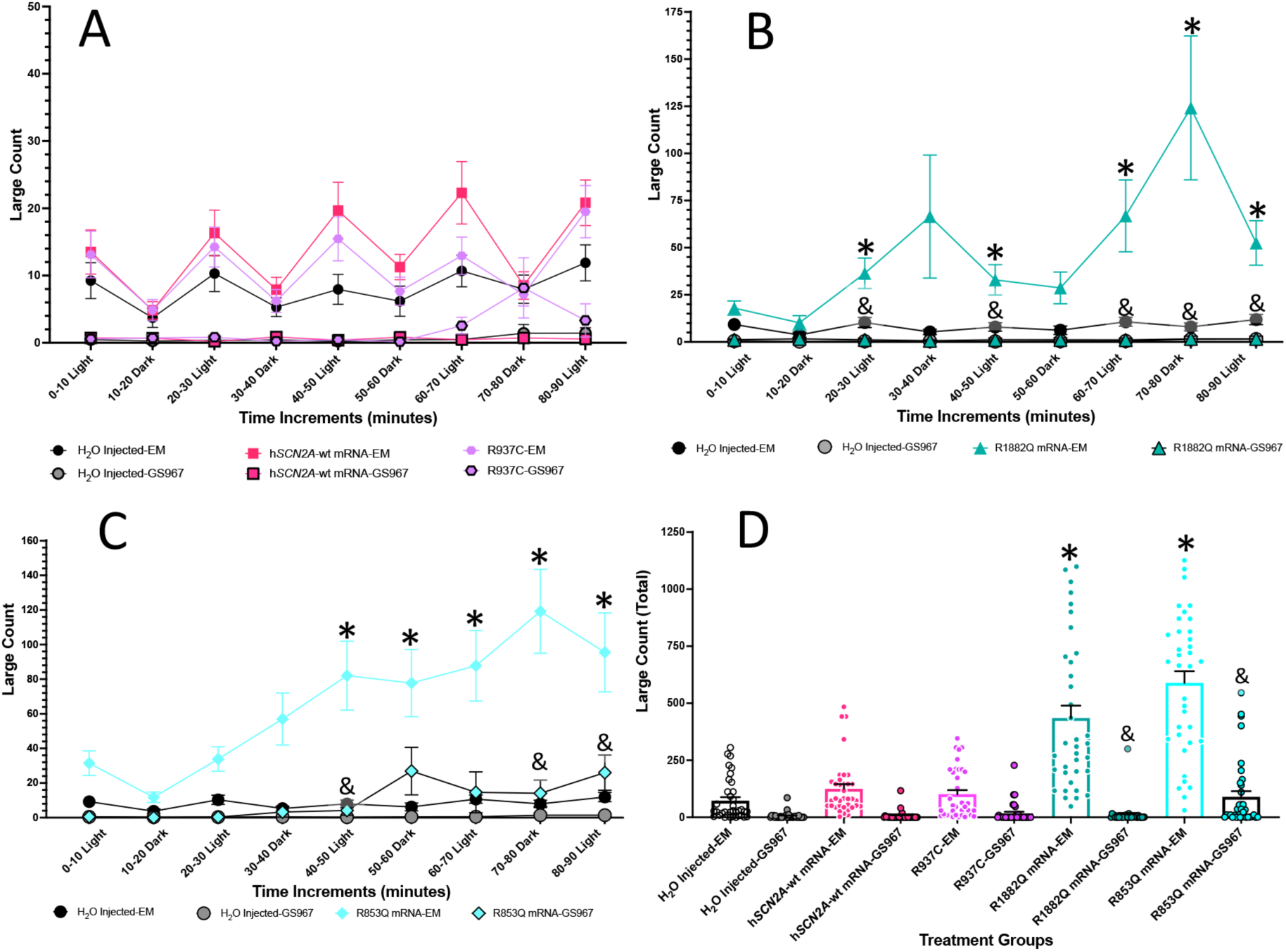
Behavioral profile and effects of GS967 on 3 dpf h*SCN2A* injected zebrafish larvae. (A-C) The average large count, number of large movement events (y-axis) of larvae is plotted against time, alternating 10-minute cycles of light and dark (x-axis) during the tracking experiment. The treatment groups are listed below the x-axis, showing the shape and color of the graph points and lines. Embryo medium (EM) is compared with ASM treatment. (A) Graph compares GS967 treatment of H_2_O injected, wild type human *SCN2A* mRNA injected, and non-seizure inducing human *SCN2A* R937C mRNA injected with untreated groups. (B) Graph compares GS967 treatment of H_2_O injected and seizure inducing human *SCN2A* R1882Q mRNA injected with untreated groups. (C) Graph compares GS967 treatment of H_2_O injected and seizure inducing human *SCN2A* R853Q mRNA injected with untreated groups. (D) A histogram representing the total large count (y-axis) summed over the 90-minute assay shows the various treatment groups. The respective treatment groups are listed below the x-axis. Statistical significance (two-way ANOVA, Tukey’s post-hoc) are marked (*=p<0.05) to indicate comparison to control, H_2_O -injected embryos. Statistical significance (two-way ANOVA, Tukey’s post-hoc) are marked (&=p<0.05) to indicate comparison to h*SCN2A*-variant vs h*SCN2A*-variant + GS967. Standard error of the mean is shown for each. n = 36 fish/treatment group. Statistics are found in the Supplementary Table S3-S4.

**Figure 4.**
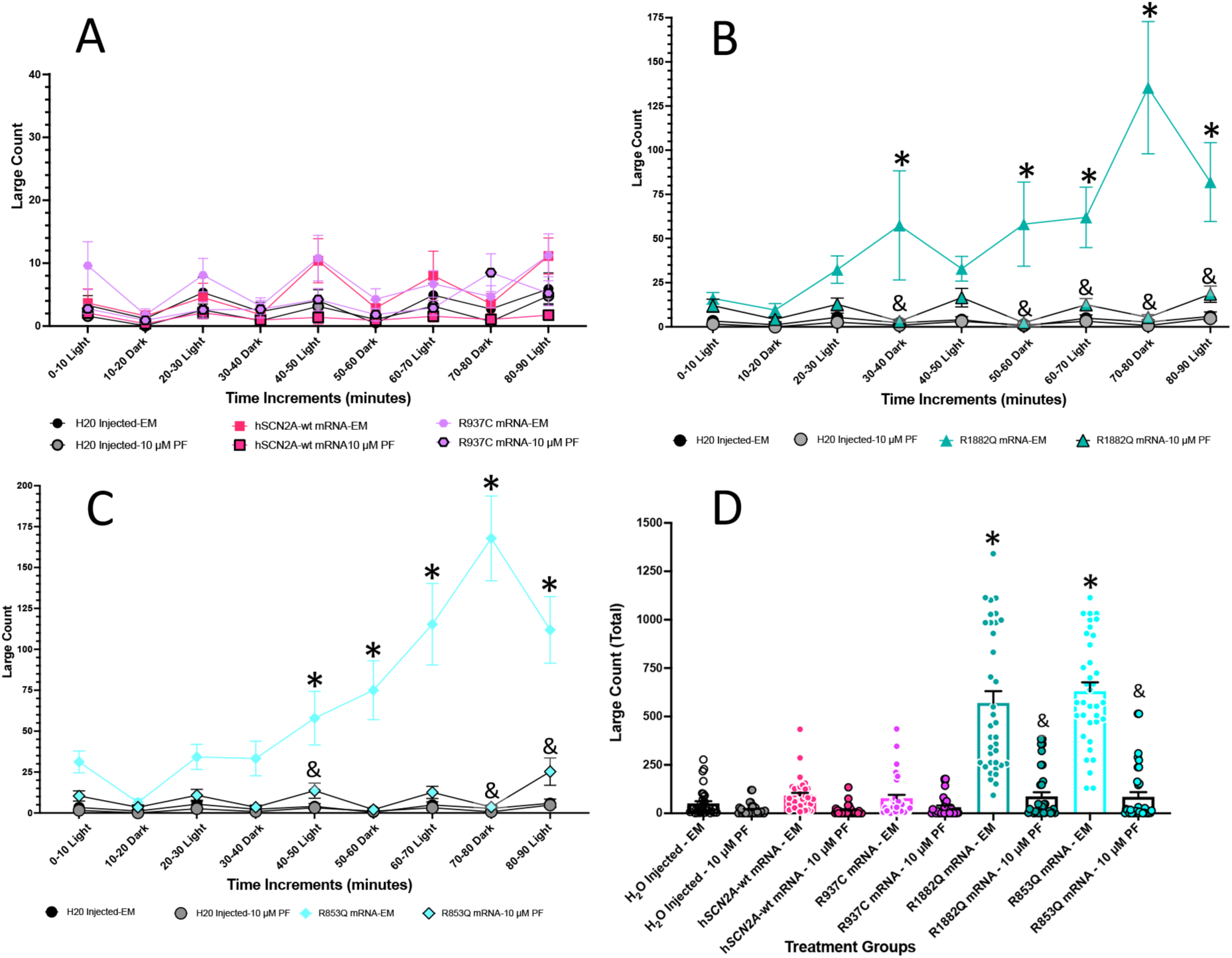
Behavioral profile and effects of PF-04856264 on 3 dpf h*SCN2A* injected zebrafish larvae. (A-C) The average large count, number of large movement events (y-axis) of larvae is plotted against time, alternating 10-minute cycles of light and dark (x-axis) during the tracking experiment. The treatment groups are listed below the x-axis, showing the shape and color of the graph points and lines. Embryo medium (EM) is compared with ASM treatment. (A) Graph compares PF-04856264 treatment of H_2_O injected, wild type human *SCN2A* mRNA injected, and non-seizure inducing human *SCN2A* R937C mRNA injected with untreated groups. (B) Graph compares PF-04856264 treatment of H_2_O injected and seizure inducing human *SCN2A* R1882Q mRNA injected with untreated groups. (C) Graph compares PF-04856264 treatment of H_2_O injected and seizure inducing human *SCN2A* R853Q mRNA injected with untreated groups. (D) A histogram representing the total large count (y-axis) summed over the 90-minute assay shows the various treatment groups. The respective treatment groups are listed below the x-axis. Statistical significance (two-way ANOVA, Tukey’s post-hoc) are marked (*=p<0.05) to indicate comparison to control, H_2_O -injected embryos. Statistical significance (two-way ANOVA, Tukey’s post-hoc) are marked (&=p<0.05) to indicate comparison to h*SCN2A*-variant vs h*SCN2A*-variant + PF-04856264. Standard error of the mean is shown for each. n = 36 fish/treatment group. Statistics are found in the Supplementary Table S5-S6.

### Behavioral profile of 3 dpf *SCN2A* injected zebrafish larvae treated with topiramate

Next, the zebrafish platform was used to evaluate the effects of ASMs on the *SCN2A* variants. TPR, GS967 and PF-04856264 treatments had no significant effect on large movement count on the H_2_O injected, *SCN2A*-wt, or R937C groups as compared to their respective untreated (EM only, no drug) groups (Fig 2A, 3A & 4A). DMSO was used to dissolve the ASMs (0.1% DMSO final concentration for all treatments). In control experiments, the 0.1% DMSO treatment did not affect activity of H_2_O injected or epileptic *SCN2A* variants (Fig 5).

**Figure 5.**
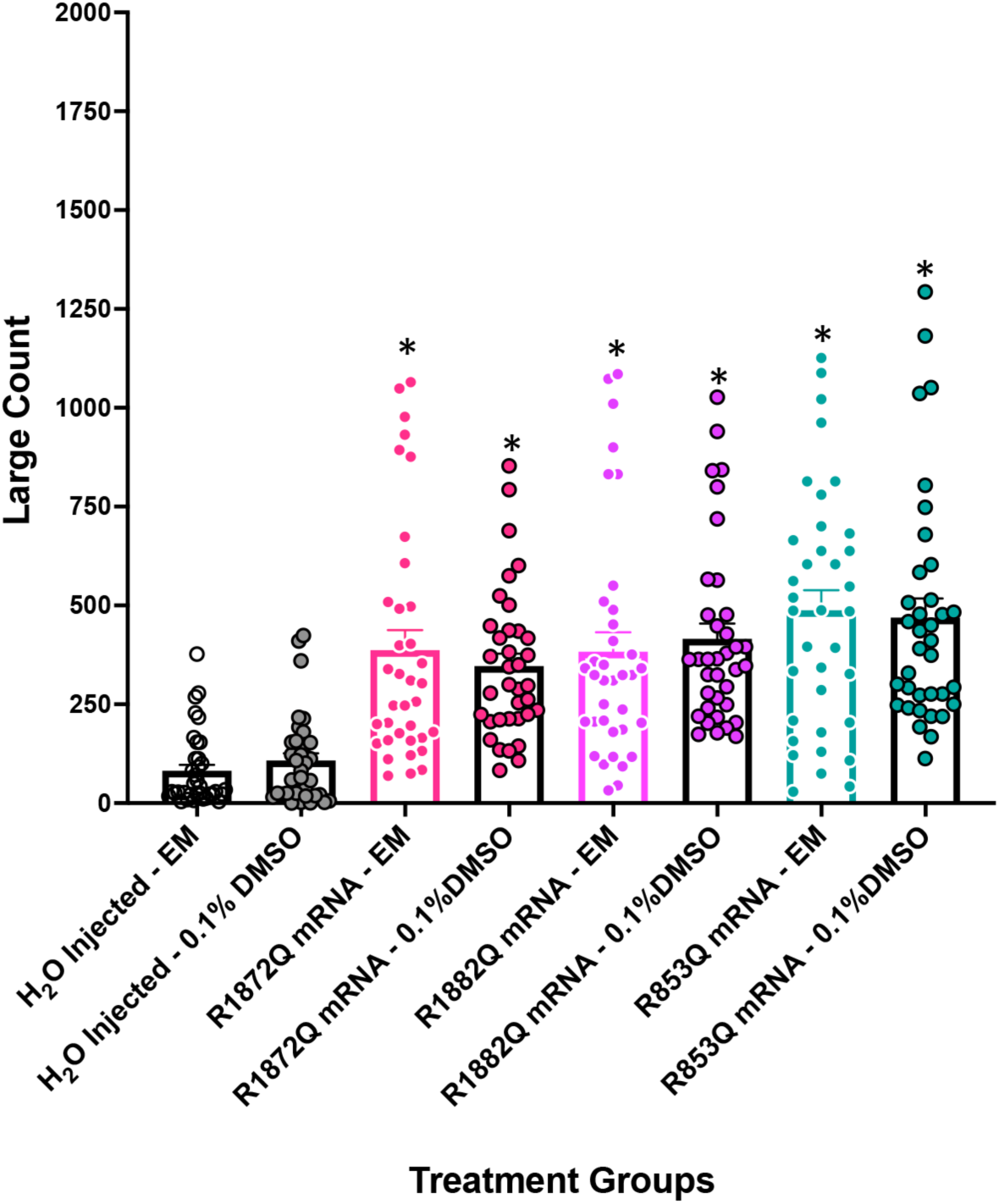
Behavioral profile and effects of 0.1% DMSO on 3 dpf h*SCN2A* and h*SCN8A* injected zebrafish larvae. The summed total large count (y-axis) over 90-minute assay using alternating light and dark periods are plotted on the histogram. The respective treatment groups are listed below the x-axis. Analyses (one-way ANOVA, Tukey’s post-hoc) are marked *(p<0.05) to indicate significant differences to H_2_O-injected embryos. There was no significant difference between the H_2_O-injected or mRNA-injected groups and their DMSO treated counterpart. Standard error of the mean is indicated on each bar. n = 36 fish/treatment group. Statistics are found in the Supplementary Table S13.

The clinically established anti-seizure compound TPR was tested first. The 200 µM TPR concentration was based on our previous experiments and pilot studies. There was significantly increased large movement counts (Fig 2B) in the R1882Q group as compared to the H_2_O group at 5 time periods (20 – 30 minutes, 40 - 50 minutes, 60 – 70 minutes, 70 - 80 minutes, 80 - 90 minutes). The R853Q group showed significant increases in large movement counts (Fig 2C) at 5 time periods (40 – 50 minutes, 50 - 60 minutes, 60 – 70 minutes, 70 - 80 minutes and 80 - 90 minutes). TPR treatment reduced large movement counts in these two *SCN2A* pathogenic variants, the R1882Q and R853Q groups, as compared to their respective untreated larvae (Fig 2B and 2C). TPR treated R1882Q injected groups showed a significant decrease in large movement counts at 5 time periods (20 – 30 minutes, 40 - 50 minutes, 60 – 70 minutes, 70 - 80 minutes, 80 - 90 minutes) as compared to untreated TPR treated R1882Q injected groups (Fig 2B). TPR treated R853Q injected showed a significant decrease in large movement counts at 3 time periods (40 – 50 minutes, 70 - 80 minutes, 80 - 90 minutes) as compared to untreated TPR treated R853Q injected groups (Fig 2C). The R1882Q and R853Q, TPR treated groups exhibited significantly lower summed total large counts over the 90 assay as compared untreated R1882Q or R853Q injected larvae (Fig 2D).

### Behavioral profile of 3 dpf *SCN2A* injected zebrafish larvae treated with GS967

An experimental ASM, GS967 was next tested. The 0.05 µM GS967 concentration was used based on our previous experiments (Milder et al., 2022) and pilot studies. There was significantly increased large movement counts in the untreated R1882Q group (Fig 3B) as compared to the H_2_O group at five time periods (20 – 30 minutes, 40 – 50 minutes, 60 – 70 minutes, 70 – 80 minutes, 80 – 90 minutes). The untreated R853Q group again showed significant increases in large movement counts (Fig 3C) as compared to the H_2_O group at five time periods (40 - 50 minutes, 50 - 60 minutes, 60 – 70 minutes, 70 - 80 minutes, 80 - 90 minutes). GS967 treatment reduced large movement counts in the *SCN2A* pathogenic variants, the R1882Q and R853Q groups as compared to their respective untreated larvae (Fig 3B-D). TPR treated R1882Q injected groups showed a significant decrease in large movement counts at five time periods (20 – 30 minutes, 40 – 50 minutes, 60 – 70 minutes, 70 – 80 minutes, 80 – 90 minutes) as compared to untreated R1882Q injected larvae. TPR treated R853Q injected groups showed a significant decrease in large movement counts in three time periods (40 - 50 minutes, 70 – 80 minutes, 80 - 90 minutes) as compared to untreated R853Q injected larvae. The R1882Q and R853Q, GS967 treated groups produced significantly lower summed total large counts over the 90 assay as compared untreated R1882Q or R853Q injected larvae (Fig 3D).

### Behavioral profile of 3 dpf *SCN2A* injected zebrafish larvae exposed to PF-04856264

The effects of PF-04856264 were examined on the zebrafish seizure activity. PF-04856264 has not been used in the zebrafish model previously. PF-04856264 is a sodium channel inhibitor with preferential activity for Na_v_1.7 but has an intermediate effect on Na_v_1.2 in electrophysiology HEK cell studies (McCormack et al., 2013; Wang et al., 2017). Dose response was measured, and 0.1 µM, 1 µM, 10 µM and 100 µM PF-04856264 reduces seizure activity. The 10 µM PF-04856264 concentration was found to exhibit a consistent effect and therefore was used in subsequent experiments described below.

As seen earlier, there was a significant increase in large movement counts (Fig 4B) in the R1882Q group compared to the H_2_O group at five time periods (30 - 40 minutes, 50 – 60 minutes, 60 – 70 minutes, 70 - 80 minutes, 80 - 90 minutes). The R853Q larvae showed significant increases in large movement counts (Fig 4C) at five time periods (40 – 50 minutes, 50 - 60 minutes, 60 – 70 minutes, 70 - 80 minutes, 80 - 90 minutes).

PF-04856264 treatment reduced large movement counts in the R1882Q and R853Q as compared to untreated groups. The 10 µM PF-04856264 treated R1882Q injected group showed significant decreases in large movement counts at five time periods (30 - 40 minutes, 50 – 60 minutes, 60 – 70 minutes, 70 - 80 minutes, 80 - 90 minutes) as compared to untreated R1882Q injected larvae (Fig 4B). The 10 µM PF-04856264 treated R853Q injected groups showed significant decreases in large movement counts at three time periods (40 – 50 minutes, 70 - 80 minutes, 80 - 90 minutes) as compared to untreated R853Q injected larvae (Fig 4C).

The total large counts summed over the 90-minute assay were compared for the various treatment groups. The R1882Q and R853Q, untreated groups produced significantly higher total large counts as compared to the H_2_O injected group (Fig 4D). The R1882Q and R853Q PF-04856264 treatment groups were significantly lower in total large counts as compared to their respective untreated group (Fig 4D).

### Pathogenic *SCN8A* variant expression in larvae induces seizure behavior

To determine whether this assay can be applied to other sodium channel genes, the R1872Q epilepsy variant of the *SCN8A* gene was tested. H_2_O injected control and wildtype *SCN8A* mRNA injected larvae did not produce seizure activity (Figs 6A, 7A & 8A). At 800 ng/μL, the R1872Q variant induced consistent seizures, producing an increase in the large count (Figs 6B, 7B & 8B). Also, the R1872Q group produced significantly higher summed total large movement counts as compared to the H_2_O injected larvae (Figs 6C, 7C & 8C).

**Figure 6.**
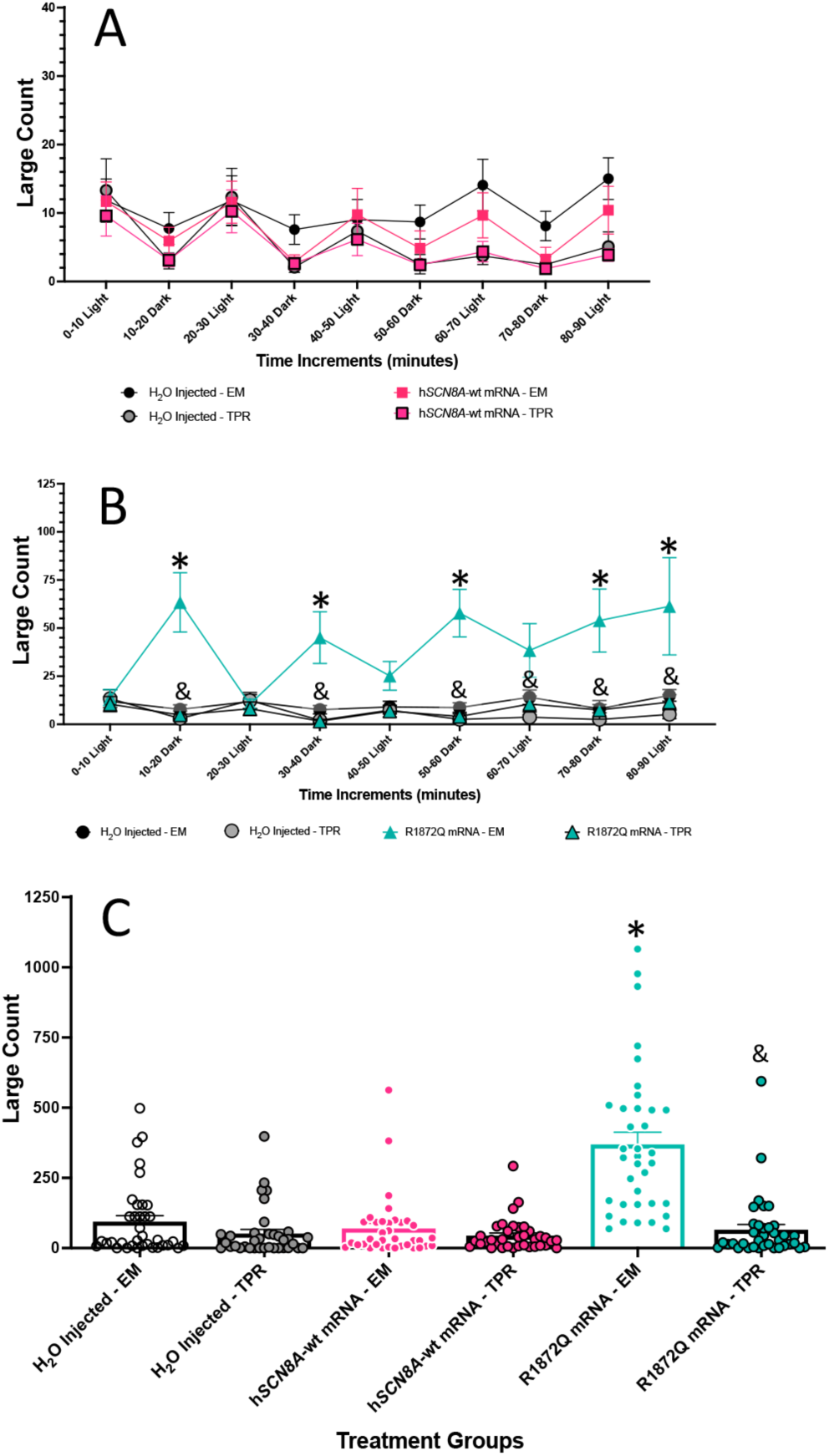
Behavioral profile and effects of topiramate on 3 dpf h*SCN8A* injected zebrafish larvae. (A and B) The average large count, number of large movement events (y-axis) of larvae is plotted against time, alternating 10-minute cycles of light and dark (x-axis) during the tracking experiment. The treatment groups are listed below the x-axis, showing the shape and color of the graph points and lines. Embryo medium (EM) is compared with ASM treatment. (A) Graph compares TPR treatment of H_2_O injected and wild type human *SCN8A* mRNA injected with untreated groups. (B) Graph compares TPR treatment of H_2_O injected and seizure inducing human *SCN8A* R1872Q mRNA injected with untreated groups. (C) A histogram representing the total large count (y-axis) summed over the 90-minute assay shows the various treatment groups. The respective treatment groups are listed below the x-axis. Statistical significance (two-way ANOVA, Tukey’s post-hoc) are marked (*=p<0.05) to indicate comparison to control, H_2_O - injected embryos. Statistical significance (two-way ANOVA, Tukey’s post-hoc) are marked (&=p<0.05) to indicate comparison to h*SCN8A*-variant vs h*SCN2A*-variant + TPR. Standard error of the mean is shown for each. n = 36 fish/treatment group. Statistics are found in the Supplementary Table S7-S8.

**Figure 7.**
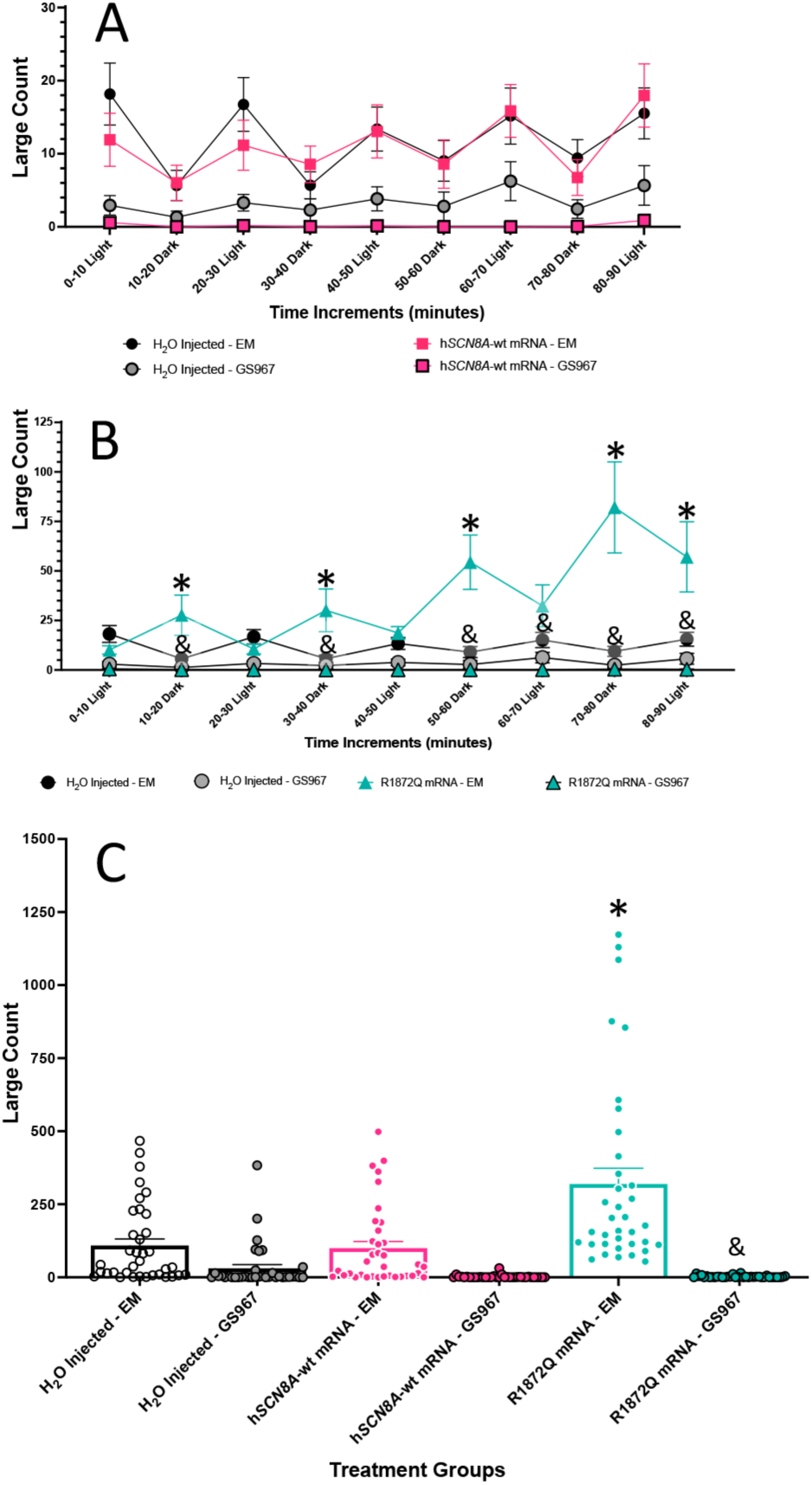
Behavioral profile and effects of GS967 on 3 dpf h*SCN8A* injected zebrafish larvae. (A and B) The average large count, number of large movement events (y-axis) of larvae is plotted against time, alternating 10-minute cycles of light and dark (x-axis) during the tracking experiment. The treatment groups are listed below the x-axis, showing the shape and color of the graph points and lines. Embryo medium (EM) is compared with ASM treatment. (A) Graph compares GS967 treatment of H_2_O injected and wild type human *SCN8A* mRNA injected with untreated groups. (B) Graph compares GS967 treatment of H_2_O injected and seizure inducing human *SCN8A* R1872Q mRNA injected with untreated groups. (C) A histogram representing the total large count (y-axis) summed over the 90-minute assay shows the various treatment groups. The respective treatment groups are listed below the x-axis. Statistical significance (two-way ANOVA, Tukey’s post-hoc) are marked (*=p<0.05) to indicate comparison to control, H_2_O - injected embryos. Statistical significance (two-way ANOVA, Tukey’s post-hoc) are marked (&=p<0.05) to indicate comparison to h*SCN8A*-variant vs h*SCN2A*-variant + GS967. Standard error of the mean is shown for each. n = 36 fish/treatment group. Statistics are found in the Supplementary Table S9-S10.

**Figure 8.**
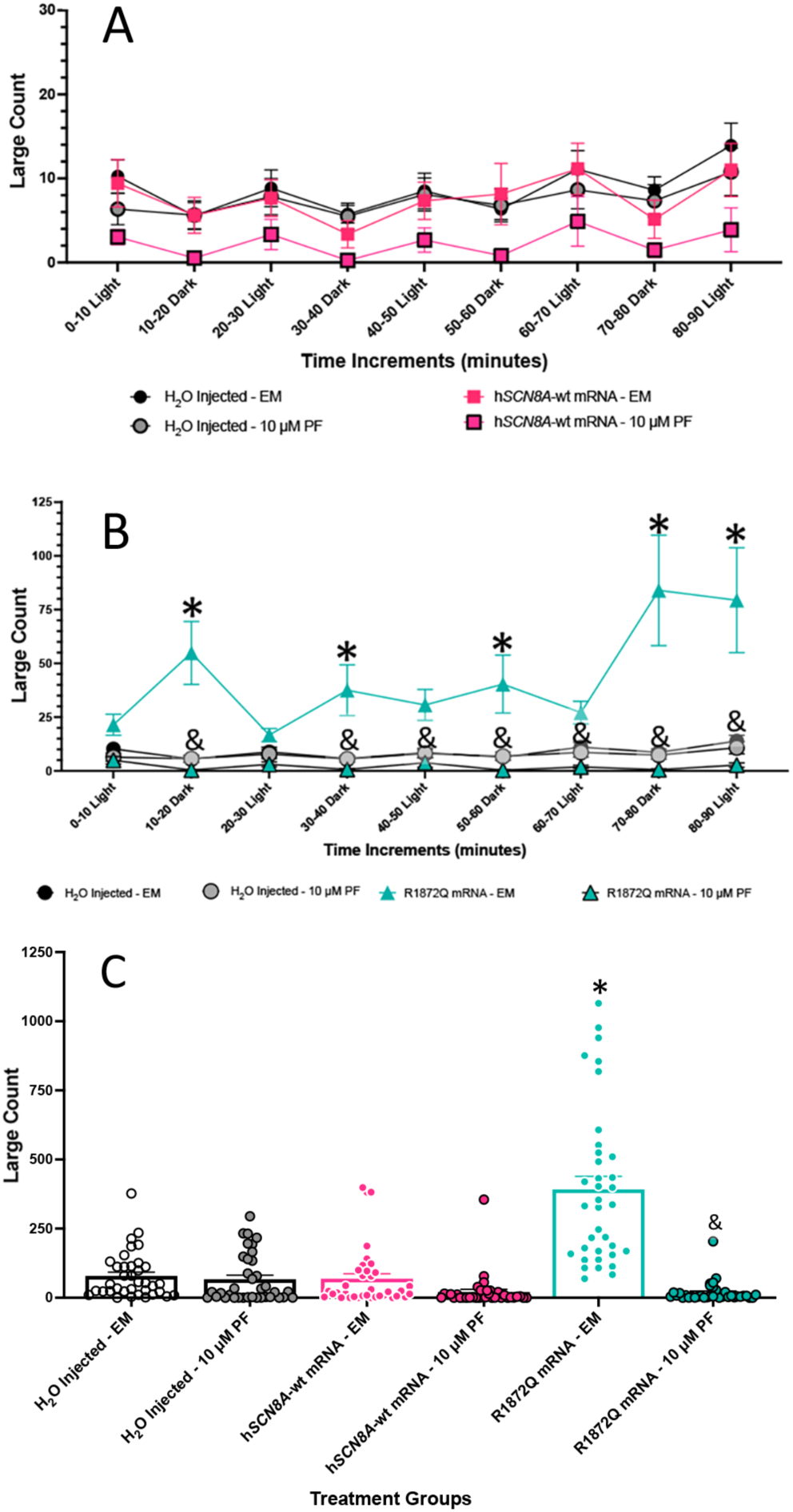
Behavioral profile and effects of PF-04856264 on 3 dpf h*SCN8A* injected zebrafish larvae. (A and B) The average large count, number of large movement events (y-axis) of larvae is plotted against time, alternating 10-minute cycles of light and dark (x-axis) during the tracking experiment. The treatment groups are listed below the x-axis, showing the shape and color of the graph points and lines. Embryo medium (EM) is compared with ASM treatment. (A) Graph compares PF-04856264 treatment of H_2_O injected and wild type human *SCN8A* mRNA injected with untreated groups. (B) Graph compares PF-04856264 treatment of H_2_O injected and seizure inducing human *SCN8A* R1872Q mRNA injected with untreated groups. (C) A histogram representing the total large count (y-axis) summed over the 90-minute assay shows the various treatment groups. The respective treatment groups are listed below the x-axis. Statistical significance (two-way ANOVA, Tukey’s post-hoc) are marked (*=p<0.05) to indicate comparison to control, H_2_O -injected embryos. Statistical significance (two-way ANOVA, Tukey’s post-hoc) are marked (&=p<0.05) to indicate comparison to h*SCN8A*-variant vs h*SCN2A*-variant + PF-04856264. Standard error of the mean is shown for each. n = 36 fish/treatment group. Statistics are found in the Supplementary Table S11-S12.

### Behavioral profile of 3 dpf *SCN8A* injected zebrafish larvae exposed to topiramate

ASMs (TPR, GS967, and PF-04856264) were tested in SCN8A assays (Figs 6, 7 & 8). The treatments of TPR, GS967, and PF-04856264 had no significant effect on large count in the H_2_O injected or *SCN8A*-wt mRNA injected groups as compared to their respective untreated groups (Figs 6A, 7A & 8A).

There was a significant increase in large movement counts in the untreated R1872Q *SCN8A* pathogenic variant group at five time periods (10 – 20 minutes, 30 - 40 minutes, 50 – 60 minutes, 70 - 80 minutes, 80 - 90 minutes) as compared to the H_2_O group (Fig 6B). TPR treatment reduced large movement counts in the R1872Q group at six time periods (10 – 20 minutes, 30 - 40 minutes, 50 – 60 minutes, 60-70 minutes, 70 - 80 minutes, 80 - 90 minutes) as compared to the untreated R1872Q larvae (Fig 6B). The R1872Q TPR treated group had significantly lower summed total large movement counts over the 90-minute assay as compared to the untreated R1872Q larvae (Fig 6C).

### Behavioral profile of 3 dpf *SCN8A* injected zebrafish larvae exposed to GS967

Again, there was a significant increase in large movement counts in the untreated R1872Q *SCN8A* pathogenic variant group at five time periods (10 – 20 minutes, 30 - 40 minutes, 50 – 60 minutes, 70 - 80 minutes, 80 - 90 minutes) as compared to the H_2_O group in the GS967 experiment (Fig 7B). GS967 treatment reduced large movement counts in the R1872Q group as compared to the untreated R1872Q larvae, showing significant decreases in large movement counts at six time periods (10 – 20 minutes, 30 - 40 minutes, 50 – 60 minutes, 60 - 70 minutes, 70 – 80 minutes, 80 - 90 minutes) as compared to the untreated R1872Q larvae (Fig 7B). The R1872Q GS967 treated group had significantly lower summed total large movement counts over the 90-minute assay as compared to the untreated R1872Q larvae (Fig 7C).

### Behavioral profile of 3 dpf *SCN8A* injected zebrafish larvae exposed to PF-04856264

Finally, there was a significant increase in large movement counts in the untreated R1872Q *SCN8A* pathogenic variant group at five time periods (10 – 20 minutes, 30 - 40 minutes, 50 – 60 minutes, 70 - 80 minutes, 80 - 90 minutes) as compared to the H_2_O group in the PF-04856264 experiment (Fig 8B). PF-04856264 treated R1872Q injected larvae had reduced large movement counts at seven time periods (10 – 20 minutes, 30 - 40 minutes, 40 - 50 minutes, 50 – 60 minutes, 60 - 70 minutes, 70 – 80 minutes, and 80 - 90 minutes) as compared to the untreated R1872Q injected larvae (Fig 8B). The PF-04856264 treated R1872Q injected group had significantly lower summed total large movement counts over the 90-minute assay as compared to the untreated R1872Q larvae (Fig 8C).

## Discussion

Epilepsy is a recurrent seizure disorder induced by abnormal electrical discharges in the brain. The causes of epilepsy include brain injury, developmental defects, tumors, and congenital syndromes. Epilepsy produces significant morbidity and mortality. Mortality usually occurs through seizure-induced accidents, but there are epilepsy syndromes that produce sudden unexpected death in epilepsy (SUDEP). Treatments for epilepsy have improved, but about 30% of epilepsy patients do not respond to known treatments.

Many pathogenic variants that produce seizure disorders in human *SCN2A* and *SCN8A* have been identified, and new variants continue to be identified. Of the known variants, few have been functionally characterized, and of those that have been characterized, many have mixed biophysical profiles. Functional characterization of these variants requires extensive experimentation, including expression in cell lines and characterization of phenotypes after expression in model organisms. Computer modeling is also being used to evaluate functional consequences of the biophysical data. However, direct evaluation of seizure activity due to the gene variants requires expression of the protein in an organism. Prior to this study, expression in whole animals was accomplished by producing stable transgenic animals. These models are very useful, but developing the models is labor intensive and time consuming.

The approach developed in this study expresses synthetic mRNAs in the zebrafish model and permits rapid phenotypic characterization of human *SCN2A* and *SCN8A* pathogenic variants. Experiments evaluated seizure inducing pathogenic variants, showing that mRNA expression of the *SCN2A*-R1882Q and *SCN8A*-R1872Q gain of function variants produced seizure-like phenotypes. The induced seizure activity was specific because expression of the wildtype mRNA or mRNA encoding a non-seizure inducing, but autism- and intellectual deficit-inducing, pathogenic variant (*SCN2A*-R937C) did not cause seizure-like behavior in the zebrafish larvae. The power of this approach is illustrated by the *SCN2A*-R853Q variant analysis. R853Q is correlated with pronounced seizure activity in patients but induces multiple loss of function changes including reduced peak current density, a depolarized voltage-dependence of activation and enhanced slow inactivation. It has also been shown to induce gating pore currents in Nav1.2 channels. Thus, our analysis indicates that the gating pore currents associated with the R853Q variant underlie a gain of function effect that induces seizure activity in intact organisms. Our new approach provides a rapid, reproducible, and specific screening tool for sodium channel genetic variants that have mixed biophysical profiles, have conflicting interpretations or are of uncertain significance.

Epilepsy induced by a pathogenic voltage-gated sodium channel variant is often resistant to current ASMs. The mRNA expression assay described here has the potential to be a rapid screening method to evaluate ASM effects on seizure-inducing pathogenic variants, which can be added to the toolbox for studying epilepsy. Seizure activity reduction was found by treating with an established ASM (topiramate) and potential ASMs (GS967 and PF-04856264). The mRNA expression assay can evaluate ASM responsiveness on patient-specific variants, providing a potential way to compare efficacies of ASMs for personalized medicine. Scale up and optimization would be necessary to ensure a rapid, reliable comparison between different ASMs. The assay could identify specific compounds or classes of compounds that are effective for reducing seizures.

The mRNA expression assay would allow comparison of related mutations in specific genes. Differences in the pattern of seizure behavior between the sodium channel variants *SCN2A* R1882Q, *SCN2A* R853Q, and *SCN8A* R1872Q were observed in our experiments. While all the three variants increased seizure activity, the time course of the seizure activity differed between these three different pathogenic variants. In future studies, the *SCN2A* R1882Q variant could be compared to other mutations at this amino acid, which include R1882L, R1882P and R1882G. The variant R1882G has a similar neutralizing amino acid change as R1882Q, and both variants show gain-of-function biophysical properties (Schwarz et al., 2016). The seizure-induction phenotype would be predicted to be similar between these variants. The R1882L and R1882P biophysical properties are unknown, and the mRNA expression assay could give insight into seizure-inducing properties. Testing additional variants that are predicted to induce gating pore currents, perhaps combined with a mutation that eliminates the central pore conductance, could provide invaluable new information on specific disease mechanisms. Overall, there is potential to correlate patient symptoms, electrophysiological findings and animal model behavior with the patient symptoms, producing an expanded dataset for analysis.

Our study was initiated due to a lack of rapid assays to evaluate seizure activity consequences for human gene variants. This zebrafish mRNA expression assay could also be used to test other seizure-inducing gain-of-function gene variants, including gene families other than voltage-gated sodium channel genes. This zebrafish mRNA expression assay could open new avenues of investigation and provide a proof-of-principle assay for epilepsy research.

Voltage-gated sodium channel function is modified by intermolecular interactions with beta-subunits of the channel, calmodulin, fibroblast growth factor homologous factors, and ankyrinG. Modifier expression variation or genetic variants could influence seizure severity in patients. This zebrafish mRNA expression assay allows for evaluation of gene variants in a consistent genetic background, controlling for background modifier expression. In addition, co-injection of mRNAs expressing seizure-inducing gene variants with mRNAs encoding these modifiers could be used to test effects of intermolecular interactions on seizure activity.

In summary, our zebrafish mRNA expression assay models seizure disorders and can be used to test ASMs. Our experiments confirmed previous findings that voltage-gated sodium channel gene variants induce seizures in animal models and allowed us to test other variants. The ASM experiments evaluated voltage-gated sodium channel variant responses to previously characterized and novel compounds. Together, our zebrafish mRNA expression assay has great potential to help understand how genetic variants produce seizures and to test ASMs on these patient specific variants, potentially leading to better treatments for the epilepsy patients.

## Acknowledgements

The Marrs Lab supported animal husbandry during our project. We thank Anna Myers and Alejandro Hernandez for technical assistance. The Marrs and Cummins Laboratory members provided helpful discussion and the sodium channel constructs used in the experiments. This work was supported by funding from the Cute Syndrome Foundation to JAM. and TRC.

## Author Contributions

Conceived and designed the experiments: PCM JAM TRC. Performed the experiments: PCM. Analyzed the data: PCM. Wrote the paper: PCM TRC JAM. Supervised JAM TRC.

## Data availability statement

All primary data are archived and can be provided upon request to the corresponding author, JAM.

## Supplementary table legends

Supplementary Table S1. Two-way ANOVA results for h*SCN2A* mRNA injected zebrafish larvae treated with topiramate.

The table displays the results of the two-way ANOVA (Fig 2a) and the resulting post-hoc Tukey comparisons and accompanying p-values for groups marked for significance within the Figure 2a graph.

Supplementary Table S2. One-way ANOVA results for h*SCN2A* mRNA injected zebrafish larvae treated with topiramate.

The table displays the results of the one-way ANOVA (Fig 2b) and the resulting post-hoc Tukey comparisons and accompanying p-values for groups marked for significance within the Figure 2b graph.

Supplementary Table S3. Two-way ANOVA results for h*SCN2A* mRNA injected zebrafish larvae treated with GS967.

The table displays the results of the two-way ANOVA (Fig 3a) and the resulting post-hoc Tukey comparisons and accompanying p-values for groups marked for significance within the Figure 3a graph.

Supplementary Table S4. One-way ANOVA results for h*SCN2A* mRNA injected zebrafish larvae treated with GS967.

The table displays the results of the one-way ANOVA (3b) and the resulting post-hoc Tukey comparisons and accompanying p-values for groups marked for significance within the Figure 3b graph.

Supplementary Table S5. Two-way ANOVA results for h*SCN2A* mRNA injected zebrafish larvae treated with PF compound.

The table displays the results of the two-way ANOVA (Fig 4a) and the resulting post-hoc Tukey comparisons and accompanying p-values for groups marked for significance within the Figure 4a graph.

Supplementary Table S6. The table displays the results of the one-way ANOVA (Fig 4b) and the resulting post-hoc Tukey comparisons and accompanying p-values for groups marked for significance within the Figure 4b graph.

Supplementary Table 7. Two-way ANOVA results for h*SCN8A* mRNA injected zebrafish larvae treated with topiramate.

The table displays the results of the two-way ANOVA (Fig 6a) and the resulting post-hoc Tukey comparisons and accompanying p-values for groups marked for significance within the Figure 6a graph.

Supplementary Table S8. The table displays the results of the one-way ANOVA (Fig 6b) and the resulting post-hoc Tukey comparisons and accompanying p-values for groups marked for significance within the Figure 6b graph.

Supplementary Table S9. Two-way ANOVA results for h*SCN8A* mRNA injected zebrafish larvae treated with GS967.

The table displays the results of the two-way ANOVA (Fig 7a) and the resulting post-hoc Tukey comparisons and accompanying p-values for groups marked for significance within the Figure 7a graph.

Supplementary Table S10. The table displays the results of the one-way ANOVA (Fig 7b) and the resulting post-hoc Tukey comparisons and accompanying p-values for groups marked for significance within the Figure 7b graph.

Supplementary Table S11. Two-way ANOVA results for h*SCN8A* mRNA injected zebrafish larvae treated with PF compound.

The table displays the results of the two-way ANOVA (Fig 8a) and the resulting post-hoc Tukey comparisons and accompanying p-values for groups marked for significance within the Figure 8a graph.

Supplementary Table S12. The table displays the results of the one-way ANOVA (Fig 8b) and the resulting post-hoc Tukey comparisons and accompanying p-values for groups marked for significance within the Figure 8b graph.

Supplementary Table S13. The table displays the results of the one-way ANOVA (Fig 5) and the resulting post-hoc Tukey comparisons and accompanying p-values for groups marked for significance within the Figure 8 graph.

## Supplemental Figure legend

Figure S1. Swimming tracings of behavioral assay voltage gated sodium channel mRNA injected zebrafish larvae.

Zebrafish larvae swimming tracing from the ZebraBox^TM^ shows effect of voltage gated sodium channel variants. For each variant or embryo medium (EM) control, a 10 minute period between 70-80 minutes of the assay in the dark. Note: Red lines illustrate large/rapid movements.

## Supplemental Movie legends

Movie S1. Video sequences of control and *hSCN2A* voltage gated sodium channel mRNA injected zebrafish larvae.

Zebrafish larvae rapid circle swimming and tonic behavior effects in seizure inducing voltage gated sodium channel variants (R1882Q and R853Q), which are not seen in controls (wildtype and R937Q).

Movie S2. Video sequences of control and *hSCN8A* voltage gated sodium channel mRNA injected zebrafish larvae.

Zebrafish larvae rapid circle swimming and tonic behavior effects in seizure inducing voltage gated sodium channel variant (R1872Q), which are not seen in control (wildtype).

## References

Afrikanova, T., A.S. Serruys, O.E. Buenafe, R. Clinckers, I. Smolders, P.A. de Witte, A.D. Crawford, and C.V. Esguerra. 2013. Validation of the zebrafish pentylenetetrazol seizure model: locomotor versus electrographic responses to antiepileptic drugs. PLoS One. 8:e54166.

Allen, N.M., J. Conroy, A. Shahwan, B. Lynch, R.G. Correa, S.D. Pena, D. McCreary, T.R. Magalhaes, S. Ennis, S.A. Lynch, and M.D. King. 2016. Unexplained early onset epileptic encephalopathy: Exome screening and phenotype expansion. Epilepsia. 57:e12–17.

Anand, G., F. Collett-White, A. Orsini, S. Thomas, S. Jayapal, N. Trump, Z. Zaiwalla, and S. Jayawant. 2016. Autosomal dominant SCN8A mutation with an unusually mild phenotype. Eur J Paediatr Neurol. 20:761–765.

Barker, B.S., M. Ottolini, J.L. Wagnon, R.M. Hollander, M.H. Meisler, and M.K. Patel. 2016. The SCN8A encephalopathy mutation p.Ile1327Val displays elevated sensitivity to the anticonvulsant phenytoin. Epilepsia. 57:1458–1466.

Ben-Shalom, R., C.M. Keeshen, K.N. Berrios, J.Y. An, S.J. Sanders, and K.J. Bender. 2017. Opposing Effects on Na(V)1.2 Function Underlie Differences Between SCN2A Variants Observed in Individuals With Autism Spectrum Disorder or Infantile Seizures. Biol Psychiatry. 82:224–232.

Berecki, G., K.B. Howell, Y.H. Deerasooriya, M.R. Cilio, M.K. Oliva, D. Kaplan, I.E. Scheffer, S.F. Berkovic, and S. Petrou. 2018. Dynamic action potential clamp predicts functional separation in mild familial and severe de novo forms of SCN2A epilepsy. Proc Natl Acad Sci U S A. 115:E5516–E5525.

Berecki, G., K.B. Howell, J. Heighway, N. Olivier, J. Rodda, I. Overmars, D.R.M. Vlaskamp, T.L. Ware, S. Ardern-Holmes, G. Lesca, M. Alber, P. Veggiotti, I.E. Scheffer, S.F. Berkovic, M. Wolff, and S. Petrou. 2022. Functional correlates of clinical phenotype and severity in recurrent SCN2A variants. Commun Biol. 5:515.

Blanchard, M.G., M.H. Willemsen, J.B. Walker, S.D. Dib-Hajj, S.G. Waxman, M.C. Jongmans, T. Kleefstra, B.P. van de Warrenburg, P. Praamstra, J. Nicolai, H.G. Yntema, R.J. Bindels, M.H. Meisler, and E.J. Kamsteeg. 2015. De novo gain-of-function and loss-of-function mutations of SCN8A in patients with intellectual disabilities and epilepsy. J Med Genet. 52:330–337.

Bromfield, E.B., J.E. Cavazos, and J.I. Sirven. 2006. An Introduction to Epilepsy. American Epilepsy Society, West Hartford CT.

Carvill, G.L., S.B. Heavin, S.C. Yendle, J.M. McMahon, B.J. O’Roak, J. Cook, A. Khan, M.O. Dorschner, M. Weaver, S. Calvert, S. Malone, G. Wallace, T. Stanley, A.M. Bye, A. Bleasel, K.B. Howell, S. Kivity, M.T. Mackay, V. Rodriguez-Casero, R. Webster, A. Korczyn, Z. Afawi, N. Zelnick, T. Lerman-Sagie, D. Lev, R.S. Moller, D. Gill, D.M. Andrade, J.L. Freeman, L.G. Sadleir, J. Shendure, S.F. Berkovic, I.E. Scheffer, and H.C. Mefford. 2013. Targeted resequencing in epileptic encephalopathies identifies de novo mutations in CHD2 and SYNGAP1. Nat Genet. 45:825–830.

Catterall, W.A. 2014. Sodium channels, inherited epilepsy, and antiepileptic drugs. Annu Rev Pharmacol Toxicol. 54:317–338.

de Kovel, C.G., M.H. Meisler, E.H. Brilstra, F.M. van Berkestijn, R. van ‘t Slot, S. van Lieshout, I.J. Nijman, J.E. O’Brien, M.F. Hammer, M. Estacion, S.G. Waxman, S.D. Dib-Hajj, and B.P. Koeleman. 2014. Characterization of a de novo SCN8A mutation in a patient with epileptic encephalopathy. Epilepsy Res. 108:1511–1518.

Do, M.T., and B.P. Bean. 2004. Sodium currents in subthalamic nucleus neurons from Nav1.6-null mice. J Neurophysiol. 92:726–733.

Epi, K.C., P. Epilepsy Phenome/Genome, A.S. Allen, S.F. Berkovic, P. Cossette, N. Delanty, D. Dlugos, E.E. Eichler, M.P. Epstein, T. Glauser, D.B. Goldstein, Y. Han, E.L. Heinzen, Y. Hitomi, K.B. Howell, M.R. Johnson, R. Kuzniecky, D.H. Lowenstein, Y.F. Lu, M.R. Madou, A.G. Marson, H.C. Mefford, S. Esmaeeli Nieh, T.J. O’Brien, R. Ottman, S. Petrovski, A. Poduri, E.K. Ruzzo, I.E. Scheffer, E.H. Sherr, C.J. Yuskaitis, B. Abou-Khalil, B.K. Alldredge, J.F. Bautista, S.F. Berkovic, A. Boro, G.D. Cascino, D. Consalvo, P. Crumrine, O. Devinsky, D. Dlugos, M.P. Epstein, M. Fiol, N.B. Fountain, J. French, D. Friedman, E.B. Geller, T. Glauser, S. Glynn, S.R. Haut, J. Hayward, S.L. Helmers, S. Joshi, A. Kanner, H.E. Kirsch, R.C. Knowlton, E.H. Kossoff, R. Kuperman, R. Kuzniecky, D.H. Lowenstein, S.M. McGuire, P.V. Motika, E.J. Novotny, R. Ottman, J.M. Paolicchi, J.M. Parent, K. Park, A. Poduri, I.E. Scheffer, R.A. Shellhaas, E.H. Sherr, J.J. Shih, R. Singh, J. Sirven, M.C. Smith, J. Sullivan, L. Lin Thio, A. Venkat, E.P. Vining, G.K. Von Allmen, J.L. Weisenberg, P. Widdess-Walsh, and M.R. Winawer. 2013. De novo mutations in epileptic encephalopathies. Nature. 501:217–221.

Estacion, M., J.E. O’Brien, A. Conravey, M.F. Hammer, S.G. Waxman, S.D. Dib-Hajj, and M.H. Meisler. 2014. A novel de novo mutation of SCN8A (Nav1.6) with enhanced channel activation in a child with epileptic encephalopathy. Neurobiol Dis. 69:117–123.

Ganguly, S., C.H. Thompson, and A.L. George, Jr. 2021. Enhanced slow inactivation contributes to dysfunction of a recurrent SCN2A mutation associated with developmental and epileptic encephalopathy. J Physiol. 599:4375–4388.

Howell, K.B., J.M. McMahon, G.L. Carvill, D. Tambunan, M.T. Mackay, V. Rodriguez-Casero, R. Webster, D. Clark, J.L. Freeman, S. Calvert, H.E. Olson, S. Mandelstam, A. Poduri, H.C. Mefford, A.S. Harvey, and I.E. Scheffer. 2015. SCN2A encephalopathy: A major cause of epilepsy of infancy with migrating focal seizures. Neurology. 85:958–966.

Hu, D., H. Barajas-Martinez, E. Burashnikov, M. Springer, Y. Wu, A. Varro, R. Pfeiffer, T.T. Koopmann, J.M. Cordeiro, A. Guerchicoff, G.D. Pollevick, and C. Antzelevitch. 2009. A mutation in the beta 3 subunit of the cardiac sodium channel associated with Brugada ECG phenotype. Circ Cardiovasc Genet. 2:270–278.

Johannesen, K.M., Y. Liu, M. Koko, C.E. Gjerulfsen, L. Sonnenberg, J. Schubert, C.D. Fenger, A. Eltokhi, M. Rannap, N.A. Koch, S. Lauxmann, J. Kruger, J. Kegele, L. Canafoglia, S. Franceschetti, T. Mayer, J. Rebstock, P. Zacher, S. Ruf, M. Alber, K. Sterbova, P. Lassuthova, M. Vlckova, J.R. Lemke, K. Platzer, I. Krey, C. Heine, D. Wieczorek, J. Kroell-Seger, C. Lund, K.M. Klein, P.Y.B. Au, J.M. Rho, A.W. Ho, S. Masnada, P. Veggiotti, L. Giordano, P. Accorsi, C.E. Hoei-Hansen, P. Striano, F. Zara, H. Verhelst, J.S. Verhoeven, H.M.H. Braakman, B. van der Zwaag, A.V.E. Harder, E. Brilstra, M. Pendziwiat, S. Lebon, M. Vaccarezza, N.M. Le, J. Christensen, S. Gronborg, S.W. Scherer, J. Howe, W. Fazeli, K.B. Howell, R. Leventer, C. Stutterd, S. Walsh, M. Gerard, B. Gerard, S. Matricardi, C.M. Bonardi, S. Sartori, A. Berger, D. Hoffman-Zacharska, M. Mastrangelo, F. Darra, A. Vollo, M.M. Motazacker, P. Lakeman, M. Nizon, C. Betzler, C. Altuzarra, R. Caume, A. Roubertie, P. Gelisse, C. Marini, R. Guerrini, F. Bilan, D. Tibussek, M. Koch-Hogrebe, M.S. Perry, S. Ichikawa, E. Dadali, A. Sharkov, I. Mishina, M. Abramov, I. Kanivets, S. Korostelev, S. Kutsev, K.E. Wain, N. Eisenhauer, M. Wagner, J.M. Savatt, K. Muller-Schluter, H. Bassan, A. Borovikov, M.C. Nassogne, A. Destree, A.S. Schoonjans, M. Meuwissen, M. Buzatu, A. Jansen, E. Scalais, S. Srivastava, W.H. Tan, H.E. Olson, T. Loddenkemper, A. Poduri, K.L. Helbig, I. Helbig, M.P. Fitzgerald, E.M. Goldberg, T. Roser, I. Borggraefe, T. Brunger, P. May, D. Lal, D. Lederer, G. Rubboli, H.O. Heyne, G. Lesca, U.B.S. Hedrich, J. Benda, E. Gardella, H. Lerche, and R.S. Moller. 2022. Genotype-phenotype correlations in SCN8A-related disorders reveal prognostic and therapeutic implications. Brain. 145:2991–3009.

Katz, E., O. Stoler, A. Scheller, Y. Khrapunsky, S. Goebbels, F. Kirchhoff, M.J. Gutnick, F. Wolf, and I.A. Fleidervish. 2018. Role of sodium channel subtype in action potential generation by neocortical pyramidal neurons. Proc Natl Acad Sci U S A. 115:E7184–E7192.

Kobayashi, Y., J. Tohyama, M. Kato, N. Akasaka, S. Magara, H. Kawashima, T. Ohashi, H. Shiraishi, M. Nakashima, H. Saitsu, and N. Matsumoto. 2016. High prevalence of genetic alterations in early-onset epileptic encephalopathies associated with infantile movement disorders. Brain Dev. 38:285–292.

Li, M., N. Jancovski, P. Jafar-Nejad, L.E. Burbano, B. Rollo, K. Richards, L. Drew, A. Sedo, J. Heighway, S. Pachernegg, A. Soriano, L. Jia, T. Blackburn, B. Roberts, A. Nemiroff, K. Dalby, S. Maljevic, C.A. Reid, F. Rigo, and S. Petrou. 2021. Antisense oligonucleotide therapy reduces seizures and extends life span in an SCN2A gain-of-function epilepsy model. J Clin Invest. 131.

Li, X., J. Zhang, X. Wu, H. Yan, Y. Zhang, R.H. He, Y.J. Tang, Y.J. He, D. Tan, X.Y. Mao, J.Y. Yin, Z.Q. Liu, H.H. Zhou, and J. Liu. 2016. Polymorphisms of ABAT, SCN2A and ALDH5A1 may affect valproic acid responses in the treatment of epilepsy in Chinese. Pharmacogenomics. 17:2007–2014.

Mason, E.R., F. Wu, R.R. Patel, Y. Xiao, S.C. Cannon, and T.R. Cummins. 2019. Resurgent and Gating Pore Currents Induced by De Novo SCN2A Epilepsy Mutations. eNeuro. 6.

McCormack, K., S. Santos, M.L. Chapman, D.S. Krafte, B.E. Marron, C.W. West, M.J. Krambis, B.M. Antonio, S.G. Zellmer, D. Printzenhoff, K.M. Padilla, Z. Lin, P.K. Wagoner, N.A. Swain, P.A. Stupple, M. de Groot, R.P. Butt, and N.A. Castle. 2013. Voltage sensor interaction site for selective small molecule inhibitors of voltage-gated sodium channels. Proc Natl Acad Sci U S A. 110:E2724–2732.

Meisler, M.H., G. Helman, M.F. Hammer, B.E. Fureman, W.D. Gaillard, A.L. Goldin, S. Hirose, A. Ishii, B.L. Kroner, C. Lossin, H.C. Mefford, J.M. Parent, M. Patel, J. Schreiber, R. Stewart, V. Whittemore, K. Wilcox, J.L. Wagnon, P.L. Pearl, A. Vanderver, and I.E. Scheffer. 2016. SCN8A encephalopathy: Research progress and prospects. Epilepsia. 57:1027–1035.

Milder, P.C., A.S. Zybura, T.R. Cummins, and J.A. Marrs. 2022. Neural Activity Correlates With Behavior Effects of Anti-Seizure Drugs Efficacy Using the Zebrafish Pentylenetetrazol Seizure Model. Front Pharmacol. 13:836573.

Nakamura, K., M. Kato, H. Osaka, S. Yamashita, E. Nakagawa, K. Haginoya, J. Tohyama, M. Okuda, T. Wada, S. Shimakawa, K. Imai, S. Takeshita, H. Ishiwata, D. Lev, T. Lerman-Sagie, D.E. Cervantes-Barragan, C.E. Villarroel, M. Ohfu, K. Writzl, B. Gnidovec Strazisar, S. Hirabayashi, D. Chitayat, D. Myles Reid, K. Nishiyama, H. Kodera, M. Nakashima, Y. Tsurusaki, N. Miyake, K. Hayasaka, N. Matsumoto, and H. Saitsu. 2013. Clinical spectrum of SCN2A mutations expanding to Ohtahara syndrome. Neurology. 81:992–998.

Novak, A.E., A.D. Taylor, R.H. Pineda, E.L. Lasda, M.A. Wright, and A.B. Ribera. 2006. Embryonic and larval expression of zebrafish voltage-gated sodium channel alpha-subunit genes. Dev Dyn. 235:1962–1973.

Ogiwara, I., K. Ito, Y. Sawaishi, H. Osaka, E. Mazaki, I. Inoue, M. Montal, T. Hashikawa, T. Shike, T. Fujiwara, Y. Inoue, M. Kaneda, and K. Yamakawa. 2009. De novo mutations of voltage-gated sodium channel alphaII gene SCN2A in intractable epilepsies. Neurology. 73:1046–1053.

Orger, M.B., and G.G. de Polavieja. 2017. Zebrafish Behavior: Opportunities and Challenges. Annu Rev Neurosci. 40:125–147.

Pan, Y., and T.R. Cummins. 2020. Distinct functional alterations in SCN8A epilepsy mutant channels. J Physiol. 598:381–401.

Patel, R.R., C. Barbosa, T. Brustovetsky, N. Brustovetsky, and T.R. Cummins. 2016. Aberrant epilepsy-associated mutant Nav1.6 sodium channel activity can be targeted with cannabidiol. Brain. 139:2164–2181.

Samanta, D., and R. Ramakrishnaiah. 2015. De novo R853Q mutation of SCN2A gene and West syndrome. Acta Neurol Belg. 115:773–776.

Sanders, S.J., A.J. Campbell, J.R. Cottrell, R.S. Moller, F.F. Wagner, A.L. Auldridge, R.A. Bernier, W.A. Catterall, W.K. Chung, J.R. Empfield, A.L. George, Jr., J.F. Hipp, O. Khwaja, E. Kiskinis, D. Lal, D. Malhotra, J.J. Millichap, T.S. Otis, S. Petrou, G. Pitt, L.F. Schust, C.M. Taylor, J. Tjernagel, J.E. Spiro, and K.J. Bender. 2018. Progress in Understanding and Treating SCN2A-Mediated Disorders. Trends Neurosci. 41:442–456.

Scharfman, H.E. 2007. The neurobiology of epilepsy. Curr Neurol Neurosci Rep. 7:348–354.

Schwarz, N., A. Hahn, T. Bast, S. Muller, H. Loffler, S. Maljevic, E. Gaily, I. Prehl, S. Biskup, T. Joensuu, A.E. Lehesjoki, B.A. Neubauer, H. Lerche, and U.B.S. Hedrich. 2016. Mutations in the sodium channel gene SCN2A cause neonatal epilepsy with late-onset episodic ataxia. J Neurol. 263:334–343.

Thompson, C.H., F. Potet, T.V. Abramova, J.M. DeKeyser, N.F. Ghabra, C.G. Vanoye, J.J. Millichap, and A.L. George. 2023. Epilepsy-associated SCN2A (NaV1.2) variants exhibit diverse and complex functional properties. J Gen Physiol. 155.

Trump, N., A. McTague, H. Brittain, A. Papandreou, E. Meyer, A. Ngoh, R. Palmer, D. Morrogh, C. Boustred, J.A. Hurst, L. Jenkins, M.A. Kurian, and R.H. Scott. 2016. Improving diagnosis and broadening the phenotypes in early-onset seizure and severe developmental delay disorders through gene panel analysis. J Med Genet. 53:310–317.

Van Wart, A., and G. Matthews. 2006. Impaired firing and cell-specific compensation in neurons lacking nav1.6 sodium channels. J Neurosci. 26:7172–7180.

Veeramah, K.R., J.E. O’Brien, M.H. Meisler, X. Cheng, S.D. Dib-Hajj, S.G. Waxman, D. Talwar, S. Girirajan, E.E. Eichler, L.L. Restifo, R.P. Erickson, and M.F. Hammer. 2012. De novo pathogenic SCN8A mutation identified by whole-genome sequencing of a family quartet affected by infantile epileptic encephalopathy and SUDEP. Am J Hum Genet. 90:502–510.

Wagnon, J.L., M.J. Korn, R. Parent, T.A. Tarpey, J.M. Jones, M.F. Hammer, G.G. Murphy, J.M. Parent, and M.H. Meisler. 2015. Convulsive seizures and SUDEP in a mouse model of SCN8A epileptic encephalopathy. Hum Mol Genet. 24:506–515.

Wang, J., S.W. Ou, and Y.J. Wang. 2017. Distribution and function of voltage-gated sodium channels in the nervous system. Channels (Austin). 11:534–554.

Westerfield, M. 2000. The zebrafish book. A guide for the laboratory use of zebrafish (Danio rerio). 4th Ed.ed. University of Oregon Press, Eugene.

Wolff, M., K.M. Johannesen, U.B.S. Hedrich, S. Masnada, G. Rubboli, E. Gardella, G. Lesca, D. Ville, M. Milh, L. Villard, A. Afenjar, S. Chantot-Bastaraud, C. Mignot, C. Lardennois, C. Nava, N. Schwarz, M. Gerard, L. Perrin, D. Doummar, S. Auvin, M.J. Miranda, M. Hempel, E. Brilstra, N. Knoers, N. Verbeek, M. van Kempen, K.P. Braun, G. Mancini, S. Biskup, K. Hortnagel, M. Docker, T. Bast, T. Loddenkemper, L. Wong-Kisiel, F.M. Baumeister, W. Fazeli, P. Striano, R. Dilena, E. Fontana, F. Zara, G. Kurlemann, J. Klepper, J.G. Thoene, D.H. Arndt, N. Deconinck, T. Schmitt-Mechelke, O. Maier, H. Muhle, B. Wical, C. Finetti, R. Bruckner, J. Pietz, G. Golla, D. Jillella, K.M. Linnet, P. Charles, U. Moog, E. Oiglane-Shlik, J.F. Mantovani, K. Park, M. Deprez, D. Lederer, S. Mary, E. Scalais, L. Selim, R. Van Coster, L. Lagae, M. Nikanorova, H. Hjalgrim, G.C. Korenke, M. Trivisano, N. Specchio, B. Ceulemans, T. Dorn, K.L. Helbig, K. Hardies, H. Stamberger, P. de Jonghe, S. Weckhuysen, J.R. Lemke, I. Krageloh-Mann, I. Helbig, G. Kluger, H. Lerche, and R.S. Moller. 2017. Genetic and phenotypic heterogeneity suggest therapeutic implications in SCN2A-related disorders. Brain. 140:1316–1336.

